# The caudate nucleus controls coordinated patterns of adaptive, context-dependent adjustments to complex decisions

**DOI:** 10.1101/568733

**Authors:** Takahiro Doi, Yunshu Fan, Joshua I. Gold, Long Ding

## Abstract

Our decisions often need to balance what we observe and what we desire. However, our understanding of how and where in the brain such decisions are made remains limited. A prime candidate for integrating sensory observations and desired rewards, and a focus of many modeling studies, is the basal ganglia pathway, which is known to make separate contributions to perceptual decisions that require the interpretation of uncertain sensory evidence and value-based decisions that select among outcome options ^1-16^. Here we report direct evidence for a causal role for a major input station of the basal ganglia, the caudate nucleus, in incorporating reward context and uncertain visual evidence to guide adaptive decision-making. In monkeys making saccadic decisions based on visual motion evidence and asymmetric reward-choice associations ^17^, single caudate neurons encoded information about both the visual evidence and the asymmetric rewards. Electrical microstimulation at caudate sites with task-modulated activity during motion viewing affected how the visual and reward information was used to form the decision. The microstimulation effects included coordinated changes in multiple computational components of the decision process, mimicking the monkeys’ voluntary adjustments in response to the asymmetric reward contexts. These results imply that the caudate nucleus plays key roles in coordinating the deliberative decision process that balances external evidence and internal preferences to guide adaptive behavior.

## Results

We trained monkeys to report their perceived motion direction of a random-dot kinematogram by making a saccadic eye movement to one of two visual choice targets at a self-determined time (Fig. 1a) ^17^. We manipulated motion strength across trials and reward-choice association across blocks of trials. The monkeys’ performance depended on both the strength and direction of the visual-motion evidence and the reward asymmetry (Fig. 1b). Likewise, single-unit activity of many caudate neurons was jointly modulated by both evidence strength and either reward context or expected reward size. For example, the activity of the example neuron depicted in Fig. 1c showed three types of modulation: 1) more activity for trials with contralateral choice, both during motion viewing and around saccade onset (Contra > Ipsi); 2) more activity during the blocks when the contralateral choice was paired with small reward and the ipsilateral choice was paired with large reward (green > purple); and 3) more activity for trials with higher coherence levels, particularly for trials with contralateral choices (dark shade > light shade). Across the population, caudate neurons showed diverse patterns of modulation by choice, reward context, expected reward size, and motion strength (Fig. 1d). A majority of neurons (101/142) showed joint modulation by sensory evidence (motion coherence corresponding to at least one of the two choices) and reward (either the lateralized reward context or reward size) in at least one epoch (Fig. 1e). Of these neurons, 50 showed such combined modulation during motion viewing, with heterogeneous modulation patterns (Extended Data Fig. 1).

**Figure 1.**
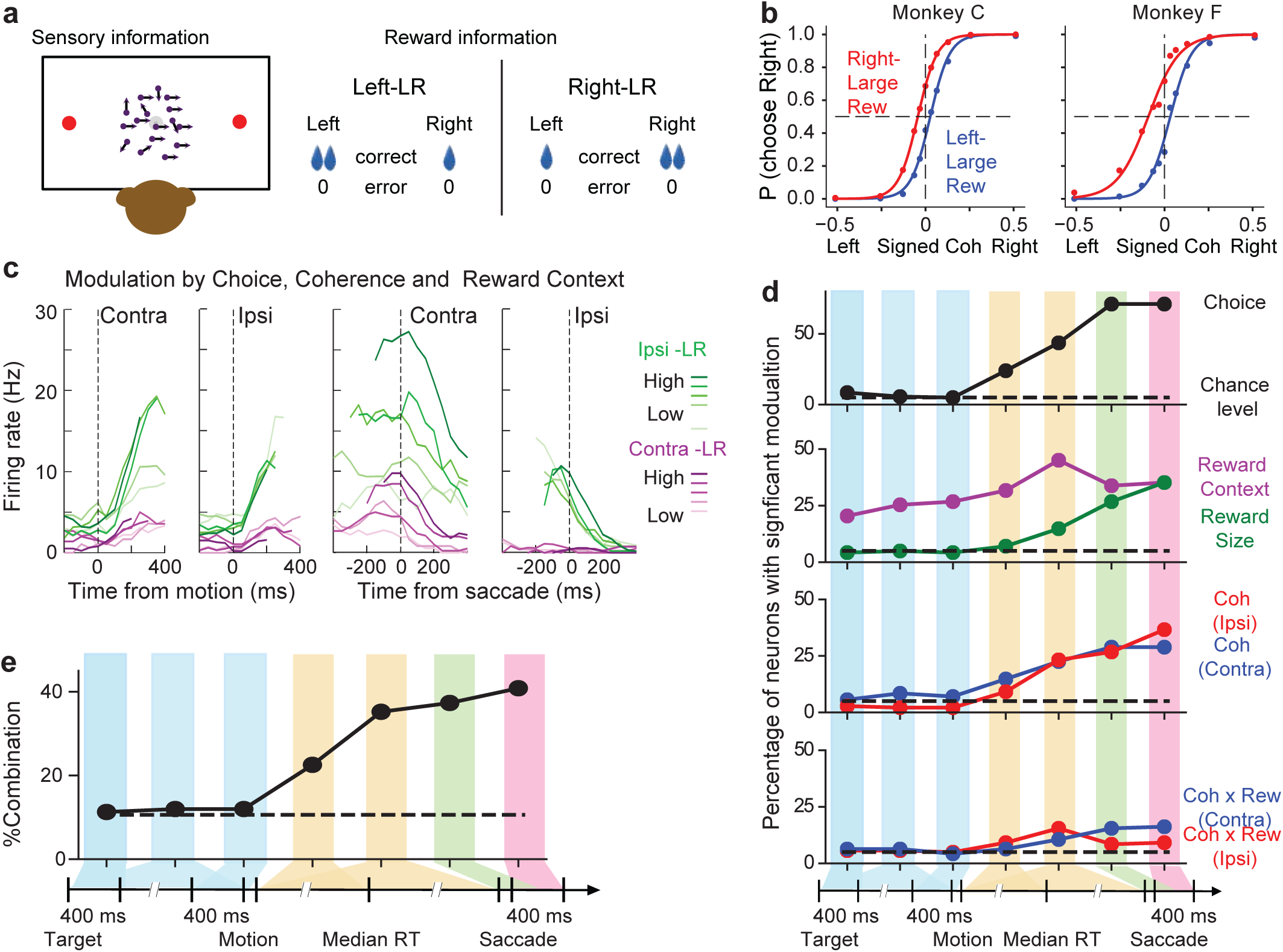
Task, behavior and caudate activity. a) For the asymmetric reward motion discrimination task, a monkey observed a random-dot motion stimulus and reported the perceived global motion with a saccade to one of two choice targets. Motion direction and strength (coherence) were pseudo-randomly selected from trial to trial. In a block of trials, a correct rightward choice was paired with a large reward and correct leftward choice was paired with a small reward. Error trials were not rewarded. The asymmetric reward contexts (left versus right) were alternated between blocks of trials. b) Average choice behavior of two monkeys (*n* = 17,493 trials from 38 sessions for monkey C, 29,599 trials from 79 sessions for monkey F). The gap between red and blue curves indicates that both monkeys were more likely to choose the choice paired with large reward. c) Example neuron showing joint modulation by choice (labels), motion strength (shading) and reward context (colors). d) Percentage of neurons showing non-zero linear regression coefficients for different regressors and task epochs (*n* = 49 for monkey C, 93 for monkey F; *t*-test, *p*<0.05). Dashed lines indicate chance level. The 7 vertical color bars indicate task epochs (defined in Methods). e) Percentage of neurons showing joint visual evidence and reward modulation. Dashed line indicates chance level.

Supporting a causal role for the caudate nucleus in decision formation, we found that electrical microstimulation during motion viewing at caudate sites with decision-related activity affected task performance (*n* = 24 sessions for monkey C, 31 for monkey F). These effects varied across sites but included choice biases (left-right shifts in the psychometric functions) that were: 1) in the same direction and of the same magnitude for the two reward contexts (Fig. 2a); 2) in opposite directions and of the same magnitude for the two reward contexts (Fig. 2b); or 3) in the same direction for the two reward contexts but of a larger magnitude for one of the two contexts (Fig. 2c). The effects of microstimulation also included changes in perceptual sensitivity (the slopes of the psychometric functions) in a reward context-dependent manner (e.g., Fig. 2b,c). Overall, microstimulation tended to induce a contralateral choice bias and a reduction in sensitivity (Fig. 2d,e, top histograms). The magnitude of these effects was larger when the contralateral choice was paired with the large reward (Fig. 2d,e, right histograms; Extended Data Fig. 2a,b). Thus, caudate neurons can causally contribute to the control of bias- and sensitivity-related computations that can be both dependent and independent of reward context.

**Figure 2.**
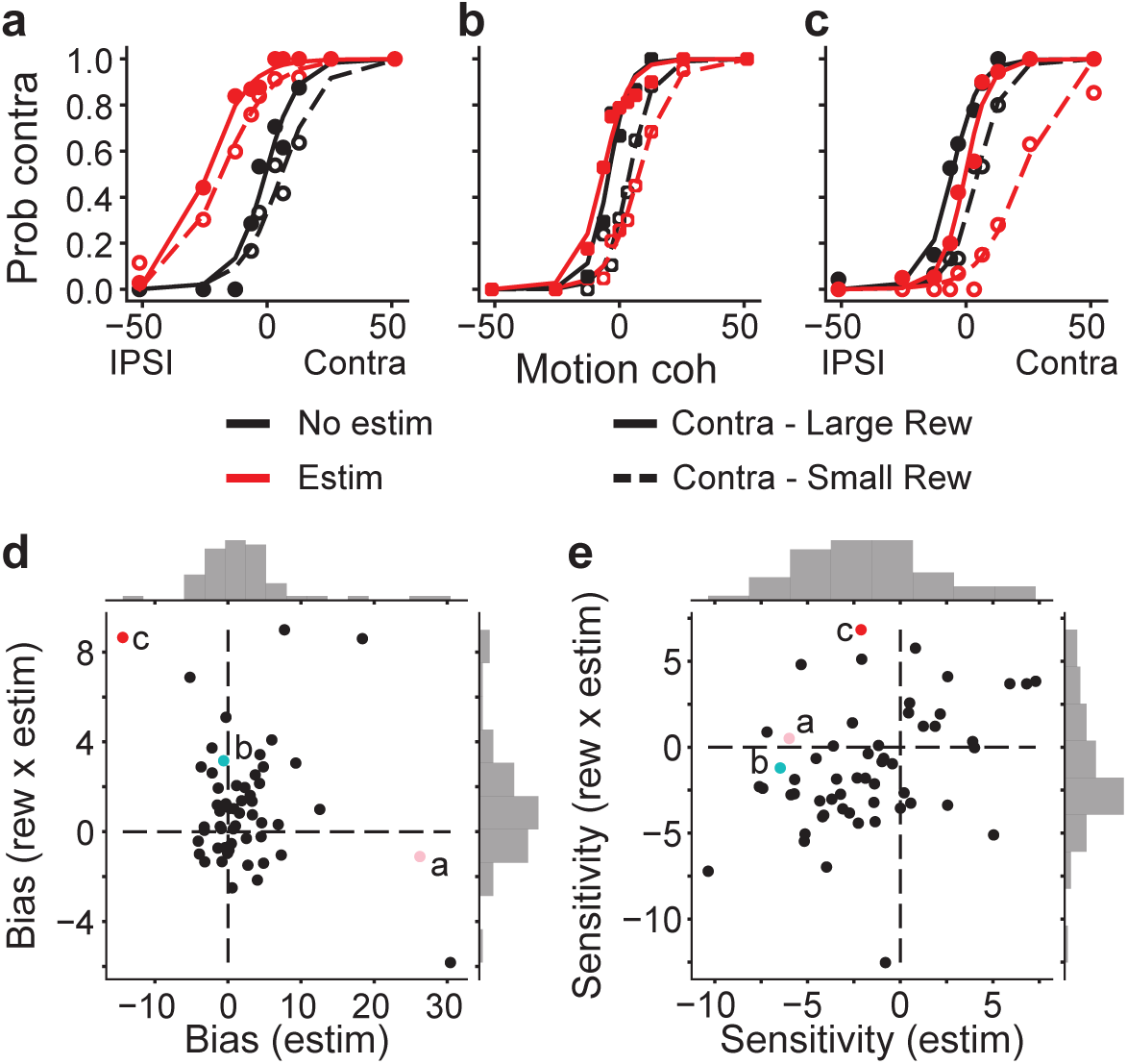
Caudate microstimulation affected monkeys’ decision behavior. a-c) Three example sessions from monkey C, showing different patterns of microstimulation effects. Black: trials without microstimulation; red: trials with microstimulation; dashed lines: logistic fits for trials in blocks when the contralateral choice was paired with small reward; solid lines: logistic fits for trials in blocks when the contralateral choice was paired with large reward. d,e) Changes in choice bias (d, logistic shift) and perceptual sensitivity (e, logistic slope) induced by electrical microstimulation (abscissa and top histograms) or interactions between electrical microstimulation and reward condition (ordinate and right histograms). d, median values=0.79%, Wilcoxon signed rank test, *p*=0.015 for the top histogram, 0.83% coherence, *p*=0.0035 for the right histogram. e, -2.1, *p*=0.0012 for the top histogram, -1.8, *p*=0.019 for the right histogram. Colored circles with labels (a, b, c) correspond to the example sessions in a–c.

We reported previously that the monkeys’ patterns of choices and response times (RTs) were well described by a drift-diffusion model (DDM; Fig. 3a) ^17-19^, in which noisy visual evidence is accumulated over time until reaching a pre-defined, time-varying, bound. In this model, the scale parameter (*k*) governs the average rate of accumulation, the bound height (*a*) governs the speed-accuracy trade-off, two parameters govern the time course of the bounds (*β_alpha* and *β_d*), and two choice-specific non-decision times account for non-perceptual processes (*t_contra* and *t_ipsi*). Using the DDM, we showed that the monkeys achieved nearly optimal rewards by: 1) a general strategy of over-biasing an offset in the neural representation of sensory information (*me*) toward the large-reward choice and compensating with asymmetric adjustments in bound heights (*z*) that favor the small-reward choice; and 2) coordinating these two kinds of adjustments to accommodate session-by-session fluctuations in the reward function (Fig. 3b)^17^.

**Figure 3.**
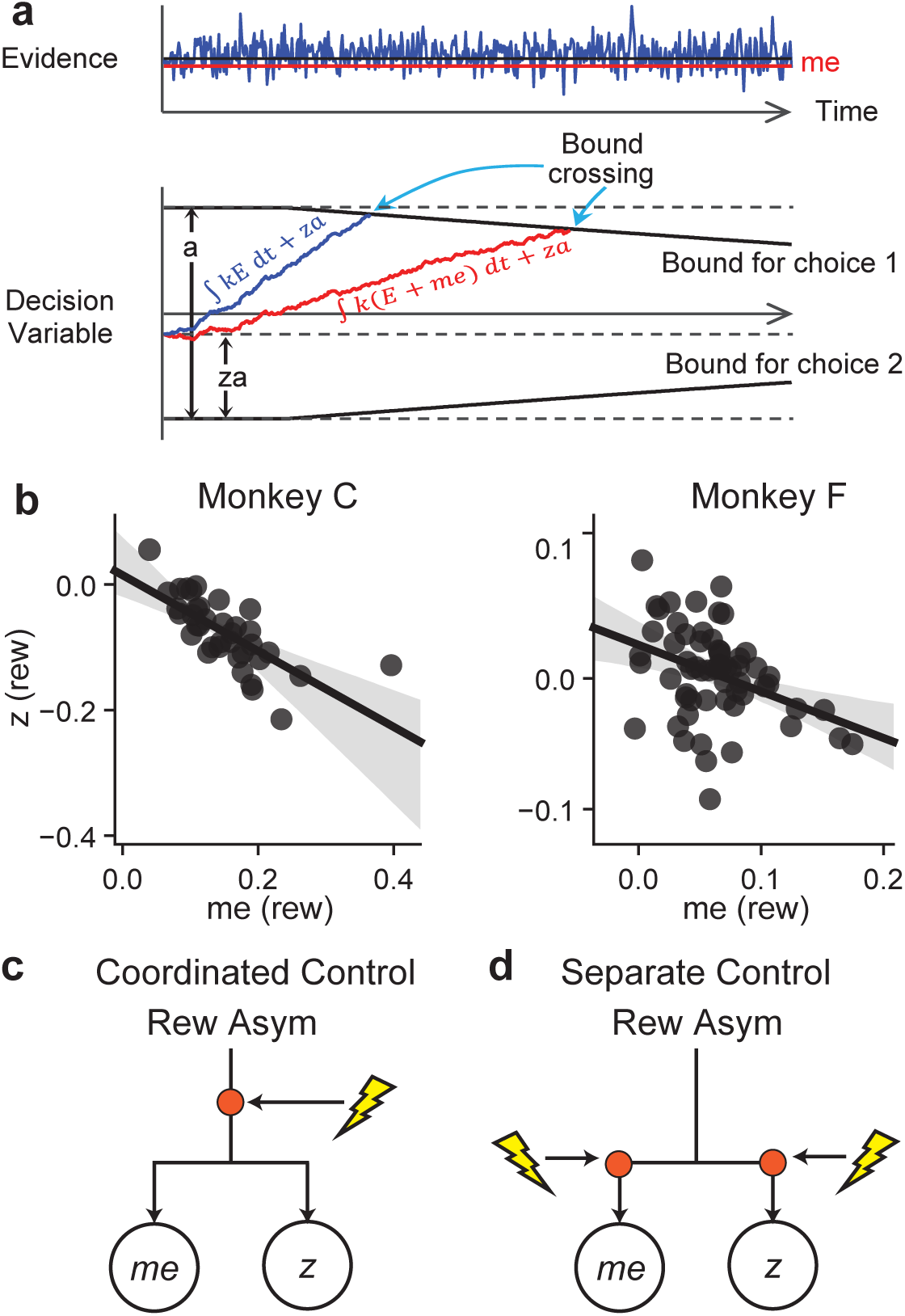
Predicted microstimulation effects on reward asymmetry-induced coordinated adjustments of multiple computational components. a) Drift-diffusion model. Motion evidence (*E*) is modeled as samples from a Gaussian distribution (mean = *k*× *signed coh*, variance = 1). The decision variable is computed as the time integral of *E* and compared at each time point to two (possibly time-varying) decision bounds. Crossing of either bound results in the corresponding choice. RT is modeled as the sum of the time to bound-crossing and a non-decision time. b) Both monkeys showed coupled adjustments in *me* and *z* parameters in the asymmetric reward contexts. Data were from a previous study without microstimulation ^17^. c,d) Alternative models of caudate’s role in coordinated decision adjustments (e.g., changes in *me* and *z* from the DDM). In principle, caudate neurons (red circles) could provide joint (c) or independent (d) control of each computational variable.

These voluntary adjustments to changes in the task conditions were affected systematically by caudate microstimulation, which evoked behavioral adjustments that were captured by similar changes in DDM parameters (Extended Data Fig. 3a-c). The effects of microstimulation, as measured via DDM fits, were variable across sessions (Extended Data Fig. 3d). However, these effects were not random but reflected a close correspondence with the session-by-session variability in the monkeys’ behavioral adjustments. In particular, for a given DDM parameter, the microstimulation effect that depended on reward context was negatively correlated across sessions with the reward asymmetry-induced adjustment (i.e., the difference in value between the two reward contexts that occurred on non-microstimulation trials; Extended Data Fig. 4a). The microstimulation effect that was shared between reward contexts was also negatively correlated with the average value between the two reward contexts on non-microstimulation trials for certain parameters (Extended Data Fig. 4b).

In principle, these microstimulation effects could reflect a role for the caudate neurons in adjusting the computational components of the decision process in a coordinated manner (Fig. 3c), separately (Fig. 3d), or both. We found evidence for both. Supporting a role for the caudate in coordinated control, the effects of microstimulation that depended on reward context reflected the monkeys’ strategy of coordinating the adjustments in multiple DDM components. For example, the negative relationship between best-fitting values of *me* and *z* in response to changes in the reward context across sessions was recapitulated by the reward context-dependent microstimulation effects on these parameters (Figs. 3b and 4a,b). Similar effects of microstimulation extended to other systematic relationships between session-specific values of pairs of DDM parameters (Fig. 4d, Extended Data Fig. 5). In contrast, the effects of microstimulation that were independent of reward context did not show similar coordination among DDM parameters (Fig. 4c,d; Extended Data Fig. 5). Coordinated effects were almost entirely absent in simulated data using DDM parameter values resampled across sessions and trial types (Extended Data Fig. 6) and thus were not an artifact of the DDM itself or the fitting procedure.

**Figure 4.**
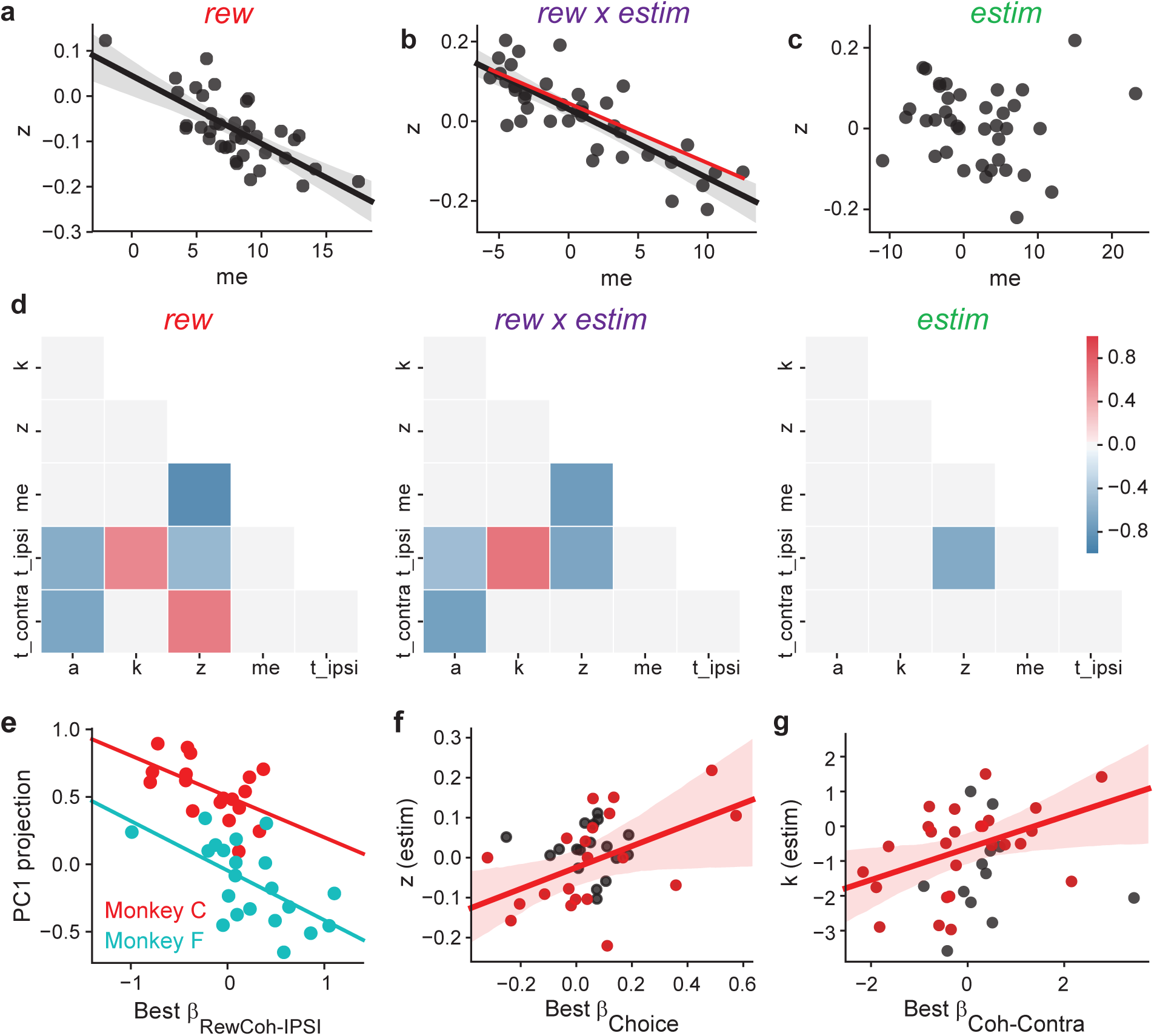
Correlated microstimulation effects on DDM parameters reflected correlated reward asymmetry effects. a–c) Scatterplots of asymmetric adjustments in momentary evidence (me) and the bound height asymmetry (z) from fits of the DDM (Fig. 3a) to behavior for the reward asymmetry-induced adjustments in non-microstimulation trials (a), reward context-dependent microstimulation effects (b), and reward context-independent microstimulation effects (c). Solid lines indicate linear regression results with significant, non-zero slope values (two-sided *t*-test, *p*<0.05). Shaded areas indicate 95% confidence intervals. The line in (a) was replotted in (b) in red for comparison. d) Pair-wise correlations between pairs of DDM parameters for the reward asymmetry-induced adjustments in non-microstimulation trials (“rew”), reward context-dependent microstimulation effects (“rew x estim”), and reward context-independent microstimulation effects (“estim”). Each color-coded box represents the average Pearson correlation coefficients of the two monkeys for changes in a pair of DDM parameters across sessions (see color bar). Pairs with non-significant correlation (*p* > 0.05, not corrected for multiple comparison) for either monkey or significant correlations but of opposite signs are shown in gray. e) Scatterplot of the regression coefficient for neural encoding of the reward size-coherence interaction for trials with ipsilateral choices and the projections of reward context-dependent microstimulation effects from the DDM fits along PC1 (see text). Red data points/line: monkey C, p=0.02; Green data points/line: monkey F, 0.004. f) Scatterplot of the regression coefficient value for neural encoding of “choice” and the reward context-independent microstimulation effect on the best-fitting value of the asymmetry bound height parameter (z). Red circles and line: sites in which the choice and reward context preferences were congruent; linear regression, p = 0.026. Black circles: sites with incongruent preferences; p = 0.72. g) Scatterplot of the regression coefficient value for neural encoding of coherence for contralateral-choice trials and the reward context-independent microstimulation effect on the best-fitting value of the scalor for evidence (k). Red circles and line: sites in which the two coefficients for Coh-Contra and Coh-Ipsi had the same signs; linear regression t-test, p = 0.023 and 0.009 for contralateral and ipsilateral choices (not shown), respectively. Black circles: sites with opposite signs; p > 0.4 for both choices.

To further examine these effects, we extracted the principal components (PCs) of the reward asymmetry-induced changes in all six DDM parameters from trials without microstimulation (Extended Data Fig. 7). These PCs describe particular patterns of coordinated adjustments to the decision process. The first three PCs captured similar structure in the reward asymmetry-induced adjustments without microstimulation (from which the PCs were extracted) and in the additional, reward context-dependent effects evoked by microstimulation: 93% and 89%, respectively, of variance explained. In contrast, the same PCs captured only 68% of variance in the microstimulation effects that did not depend on reward context. These results are consistent with roles for the caudate in adjusting the computational components of the decision process in both a coordinated manner, for changes that did depend on reward context, and separately, for changes that did not depend on reward context.

These microstimulation effects also depended on certain properties of neural selectivity at the sites of microstimulation. For example, we used the first PC from the reward asymmetry-induced changes as a proxy for the dominant coordination pattern and projected the effects of microstimulation that depended on reward context onto this PC. Such a projection was negatively correlated with the strength of neural modulation by the interaction between reward size and motion strength (Fig. 4e). This result implies that neurons that combine reward and visual information appear to play a particularly direct role in coordinating the patterns of adjustments used in these decisions. Two other effects were more consistent with a role for the caudate in adjusting components of the decision process separately. First, the strength of neural selectivity for choice was correlated with the microstimulation effect on the asymmetric bound height (*z*), but only when choice and reward context modulations were congruent (i.e., higher activity for the ipsi-/contra-lateral choice and when the ipsi-/contra-lateral choice was paired with large reward; Fig. 4f). Second, the strength of neural modulation by coherence was correlated with the effect on the average rate of evidence accumulation (*k*), but only when the modulation had the same sign for both choices (Fig. 4g). These results imply that separate subsets of caudate neurons may help control choice biases towards a particular alternative ^11, 20, 21^ and the monkeys’ perceptual sensitivity, respectively.

Collectively, our results suggest that caudate neurons causally contribute to deliberative decisions that combine reward-asymmetry and visual information. Their contributions include integrating these two sources of information at the level of single neurons and across the population of neurons, driving coordinated adjustments of multiple components of decision-related computations to rationally balance these sources of information, and mediating more directly the implementation of these computational components. The caudate nucleus’ role in coordinating decision-related computations is reminiscent of the role of premotor areas in controlling coordinated movements ^22^ and may reflect a general organization principle in which motor or cognitive primitives are aggregated into behaviorally relevant combinations. These complex and flexible contributions might also partially explain the complicated nature of decision-making impairments with striatal dysfunction, such as in addiction ^23^.

## Methods

We used two adult male rhesus macaques (*Macaca mulatta*) for this study. They were first trained extensively on an equal-reward reaction-time random-dot motion discrimination task ^3, 24, 25^ and then trained with the asymmetric-reward contexts ^17^. All training and experimental procedures were in accordance with the National Institutes of Health Guide for the Care and Use of Laboratory Animals and were approved by the University of Pennsylvania Institutional Animal Care and Use Committee.

### Behavioral task

Task details are reported elsewhere ^17^. Briefly, a trial began with a central fixation point presentation. Upon acquiring and maintaining fixation, two choice targets were presented to inform the monkeys the two possible motion directions. After a random delay picked from a truncated exponential distribution (mean = 0.7 s, range: 0.4-2.5 s), the fixation point was dimmed and a random-dot kinematogram was shown at the center of the screen (“dots onset”). For each trial, the kinematogram had a constant speed of 6°/s, aperture size of 5°, and randomly interleaved motion direction and strength (five levels of coherence: 3.2, 6.4, 12.8, 25.6, 51.2%). The monkey reported the perceived motion direction by making a self-timed saccade to the corresponding choice target. A minimum 50-ms latency was imposed, although the monkeys rarely made fast-guess responses during this study. Once the monkey’s gaze exited the fixation window (4° square window), the kinematogram was extinguished. Once the monkey’s gaze reached the choice target window (4° square window), a 400-ms minimum fixation time was imposed to register the monkey’s choice. Correct choices were rewarded with juice. Error choices were not rewarded and penalized with a timeout before the next trial (3 s for monkey F, 0.5–2 s for monkey C).

Two asymmetric reward contexts were alternated in a block design. In Contra-LR blocks, the choice contralateral to the recording/stimulation site was paired with large reward. In Ipsi-LR blocks, the choice ipsilateral to the recording/stimulation site was paired with large reward. The other choice was paired with small reward. At the start of each block, the choice targets were presented with different colors to signal the current reward context to the monkeys, followed by two additional high-coherence trials to allow the monkeys to experience the current reward context. These trials were excluded from analysis.

### Data acquisition

Eye position was monitored using a video-based system (ASL) sampled at 240 Hz. Response time (RT) was measured as the time from stimulus onset to saccade onset, the latter identified offline with respect to velocity (> 40°/s) and acceleration (> 8000°/s2). Single-unit recordings focused on putative project neurons ^24^. We searched for task-relevant neurons while the monkeys performed the equal-reward motion discrimination task with horizontal dots motions and determined the presence of task-related modulation of neural activity by visual and audio inspection of ∼10–20 trials. For analyses of neural response properties, only well-isolated single units were included. For analyses of microstimulation effects, sites with either single- or multi-unit task-related modulations were used. Neural signals were amplified, filtered and stored using a MAP acquisition system (Plexon, Inc.), along with time-stamped event codes, analog eye position signals and trial parameter values. Single unit activity was identified by offline spike sorting (Offline Sorter, Plexon, Inc.). Multiunit activity was measured using waveforms that passed an offline amplitude threshold. For the microstimulation experiments, we first identified a caudate site with task-related activity and then interleaved trials with and without microstimulation at a 1:1 ratio. Electrical microstimulation was delivered during motion stimulus presentation (negative-leading bipolar current pulses, 300 Hz, 50 μA, 250 μs pulse duration) in half of trials randomly interleaved with no-stimulation trials. Caudate microstimulation with these parameters do not evoke saccades ^3, 26-28^.

### Neural data analysis

For each single/multi-unit dataset, we computed the average firing rates in seven task epochs: three epochs before motion stimulus onset (400 ms window beginning at target onset, variable window from target onset to dots onset, and 400 ms window ending at dots onset), two epochs during motion viewing (a fixed window from 100 ms after dots onset to 100 ms before median RT and a variable window from 100 ms after dots onset to 100 ms before saccade onset), a peri- saccade 300 ms window beginning at 100 ms before saccade onset, and a post-saccade 400 ms window beginning at saccade onset (before reward delivery). For each unit, a multiple linear regression was performed for each task epoch separately.

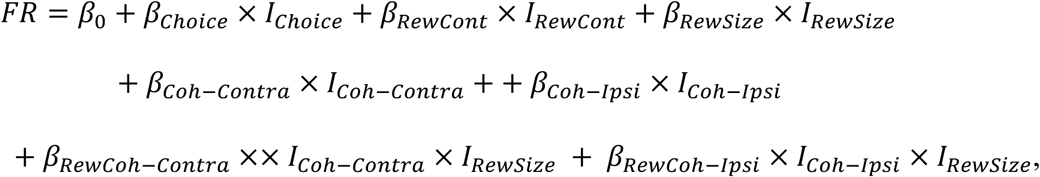

where 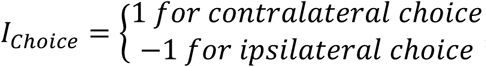,

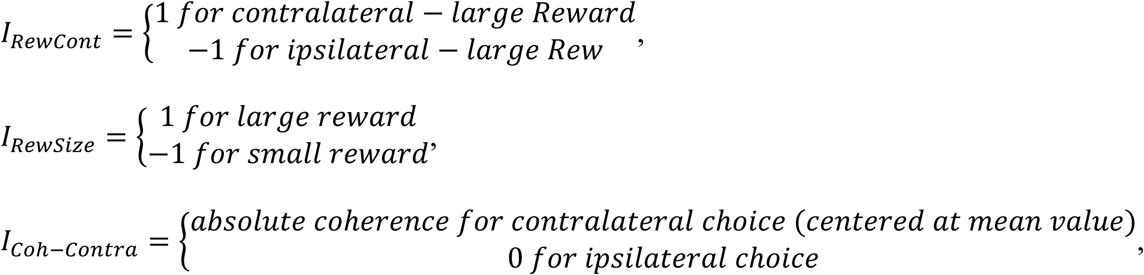

and 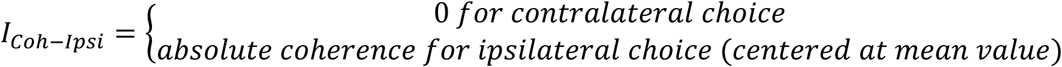.

Significance of non-zero coefficients was tested using *t*-test (criterion: *p* = 0.05).

For the microstimulation experiments, we recorded single- or multi-unit activity before microstimulation and performed multiple linear regression as above for each unit separately and normalized the regression coefficients to *β*_0_. If more than one unit was recorded at a site, for each regressor, we used the beta value associated with the lowest *p* value.

### Behavioral analysis

For each microstimulation session, a logistic function was fitted to the choice data for all trials:

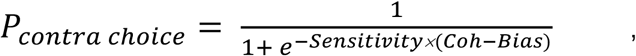

Where

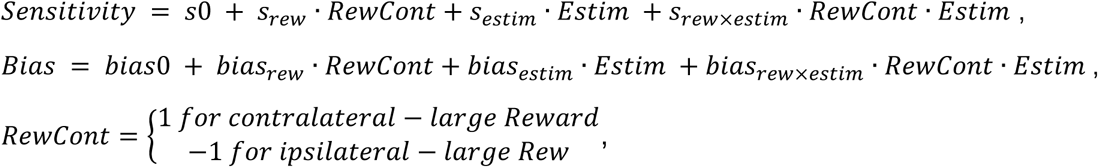

and 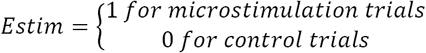.

We fitted the choice and RT data to different variants of the drift-diffusion model (DDM; Fig. 3a and Extended Data Fig. 3). The basic DDM assumed that the decision variable (DV) is the time integral of evidence (E), which was modeled as a Gaussian distributed random variable,

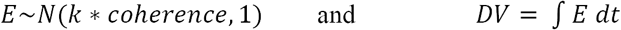

The scale parameter *k* controlled the drift rate. At each time point, the DV was compared with two collapsing choice bounds ^19^. The time course of the choice bounds was specified as *a*/(1 + *e*^*β*_*alpha*(*t*-*β*_*d*)^), where *β*_*alpha* and *β*_*d* controlled the rate and onset of decay, respectively. If DV crossed the upper bound first, a contralateral choice was made; if DV crossed the lower bound first, an ipsilateral choice was made. RT was modeled as the sum of the time till first bound crossing and saccade-specific non-decision times that accounted for evidence-independent sensory/motor delays. Two types of biases were used to account for reward asymmetry-induced biases, a bias in drift rate (*me*) and a bias in the starting point (*z*)^17^.

DDM model fitting was performed, separately for each session, using the maximum *a posteriori* estimate method (python v3.5.1, pymc 2.3.6) and prior distributions suitable for human and monkey subjects ^29^. We performed at least five runs for each variant and used the run with the highest likelihood for further analyses. As a sanity check for the quality of fits, we compared the perceptual sensitivity and choice bias estimated from logistic function fits to those estimated using the “NoCollapse” variant. Both perceptual sensitivity and choice bias were highly correlated between these two estimation methods (Pearson correlation coefficients = 0.85 and 0.98, respectively; *p* < 1e-50 for both).

We used eight variants of the DDM model: in the “Full” model, all parameters were allowed to vary by reward context and microstimulation status; in the “NoEstim” model, all parameters were allowed to vary by reward context, but not microstimulation status (Extended Data Fig. 3a); in the “NoCollapse” model, *β*_*alpha* and *β*_*d* were fixed across microstimulation status; in the “NoA”, “NoK”, “NoME”, and “NoZ” models, *a, k, me*, and *z* were fixed across microstimulation status, respectively; and in the “NoT0” model, saccade-specific non-decision times (*t_contra* and *t_ipsi*) were fixed across microstimulation status (Extended Data Fig. 3b,c). We used the Akaike information criterion (AIC) for model comparisons, with lower values indicating more parsimonious model variants.

For a given DDM parameter, we parsed the different effects as follows (LR: large reward):

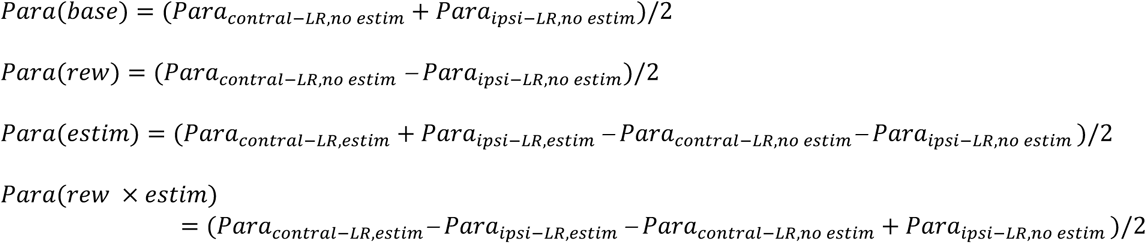

To control for potential artifacts from our analysis methods, we pooled the fitted values across reward-microstimulation conditions and sessions for each parameter, resampled from these values with replacement for each session and reward-microstimulation condition, simulated sessions with matched numbers of trials for each condition using the resampled parameter values, and re-fitted these simulated data with the Full and NoCollapse DDM models. The fitted values were used to compute the correlation patterns shown in Extended Data Fig. 6.

Principal component analysis (PCA) in Extended Data Fig. 7 was performed using python scikit-learn v0.18.1. Before PCA, the DDM parameters were pooled across “rew”, “rew x estim” and “estim” measurements to find the range for each parameter. The DDM parameters were then normalized linearly to these ranges such that each normalized parameter was in the range of [0,1], but not mean-subtracted. For data presented in Fig. 4e, the microstimulation effects that depended on reward context was projected onto the first PC extracted from the reward asymmetry-induced adjustments in non-microstimulation trials.

## Acknowledgments

We thank Jean Zweigle for animal care. This work was supported by NIH National Eye Institute (R01-EY022411) (L.D. and J.I.G), University of Pennsylvania (University Research Foundation Pilot Award) (L.D.), and Hearst Foundations Graduate student fellowship (Y.F.).

## Author contributions

Conceptualization, L.D. and J.I.G.; Methodology, Investigation, all authors; Original Draft, L.D; Writing – Review and Editing, all authors.; Supervision and Project Administration, L.D.; Funding Acquisition, L.D. and J.I.G.

## The authors declare no competing interests

## Data availability

The datasets generated during and/or analyzed during the current study are available from the corresponding author on reasonable request.

---

**Extended Data Figure 1.**
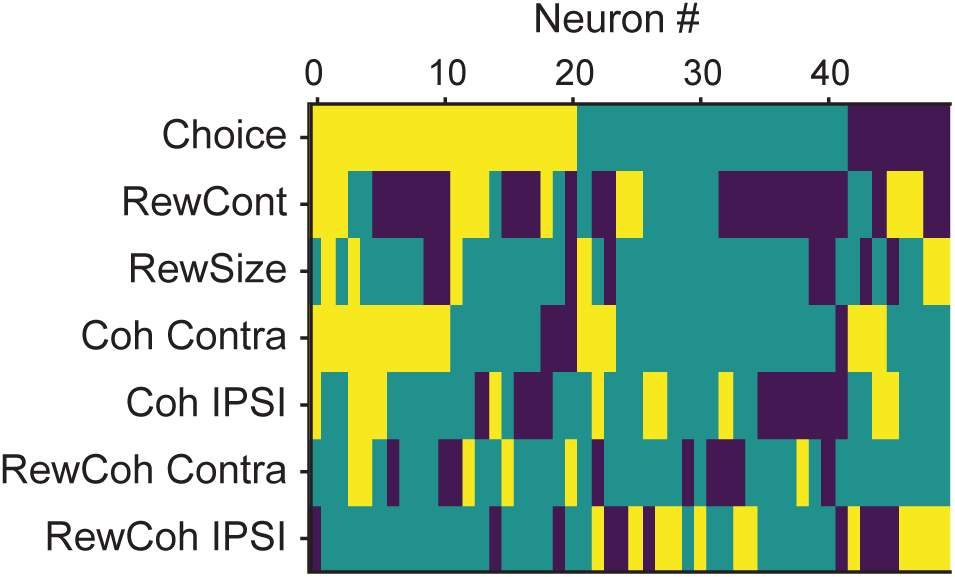
Modulation patterns of “combination neurons” during motion viewing. Color map showing the presence of significant non-zero regression coefficients for each combination neuron (in rows) during the epoch from 100 ms after motion onset until 100 ms before saccade onset. Yellow: positive; dark blue: negative; green: not significant; criterion: *p* = 0.05. Neurons were sorted by the signs of the coefficients.

**Extended Data Figure 2.**
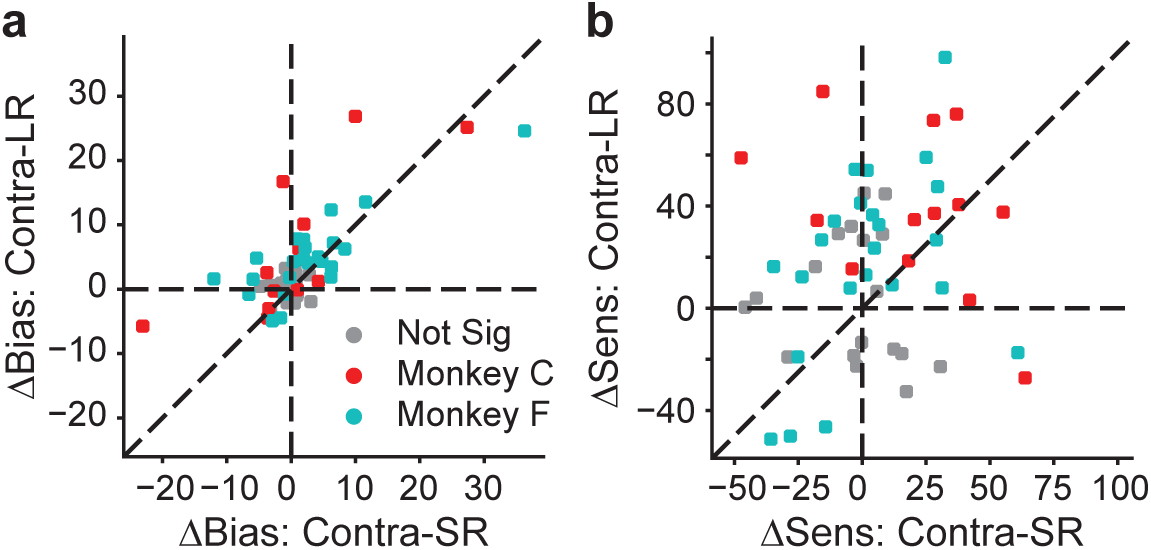
Scatterplots of effects of microstimulation on choice bias (a) and perceptual sensitivity (b) for the two reward contexts (contralateral-small-reward blocks on the abscissa, contralateral-large-reward blocks on the ordinate), measured from logistic fits to choice data. Gray data points represent sessions without a significant effect for either reward context (logistic regression, *p*>0.05).

**Extended Data Figure 3.**
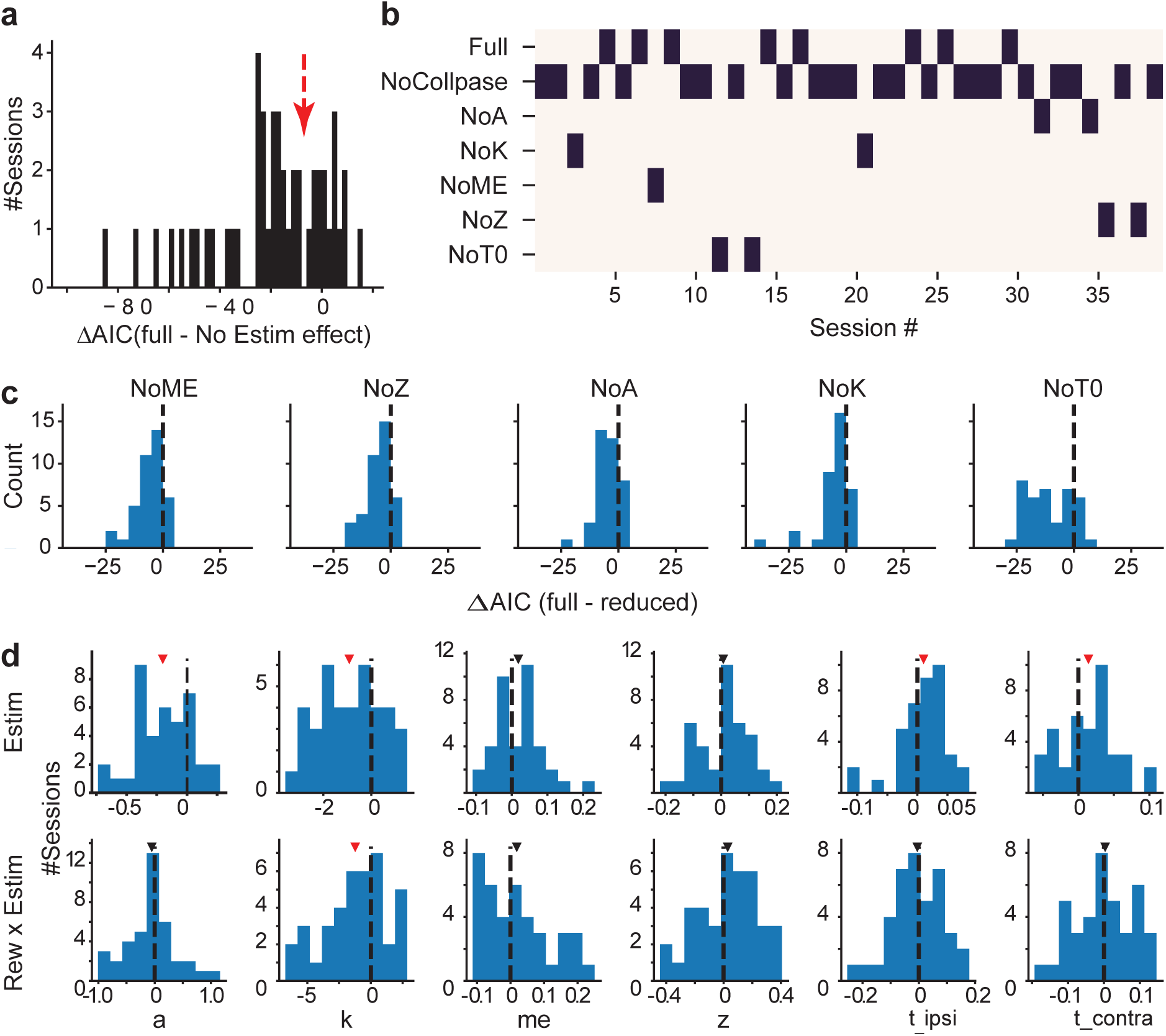
DDM fitting results. a) Histogram of the difference in AIC between the full model, in which all DDM parameters were allowed to vary by reward context and microstimulation status, and a reduced model, in which all DDM parameters were allowed to vary by reward context but not microstimulation status. Negative AIC implies that the full model is better. The red arrow indicates the criterion we used and corresponds to the gap in the histogram. b) Map showing the best DDM variant (lowest AIC, black bar) for sessions that showed an effect of microstimulation, which was defined as having a smaller AIC with the full model than a reduced model without microstimulation-induced changes (39 sessions to the left of the red arrow in a). See Methods for model variant definitions. c) Histograms of differences in AIC between the full model and reduced models. Mean ΔAIC values were negative for all reduced models (*t*-test, *p* < 0.001 for all). d) Histograms of best-fitting DDM parameters from the best model for each session. The top row (“Estim”) shows reward context-independent microstimulation effects. The bottom row (“Rew x Estim”) shows reward context-dependent microstimulation effects. Triangles indicate median values; red triangles indicate non-zero median values (Wilcoxon signed-rank test, *p* < 0.05, not corrected for multiple comparisons).

**Extended Data Figure 4.**
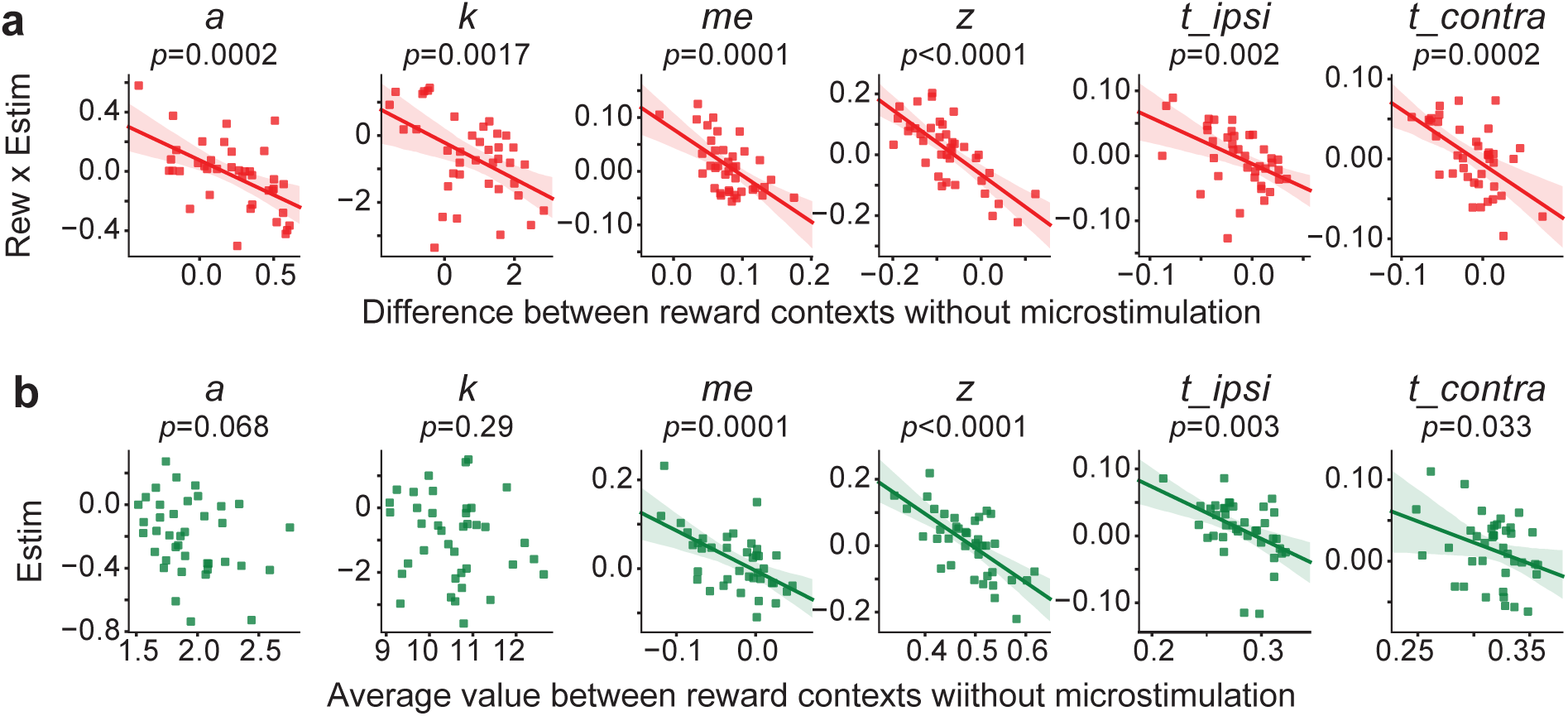
Relationships between microstimulation effects and monkeys’ voluntary adjustments. a) Scatterplots of the effects of microstimulation that depended on reward context (ordinate) and reward asymmetry-induced adjustments without microstimulation (abscissa). Lines: linear regressions. P values were from *t-*test. b) Scatterplots of the average effects of microstimulation between reward contexts (ordinate) and average adjustments between reward contexts without microstimulation (abscissa). Lines: linear regressions. P values were from *t-*test.

**Extended Data Figure 5.**
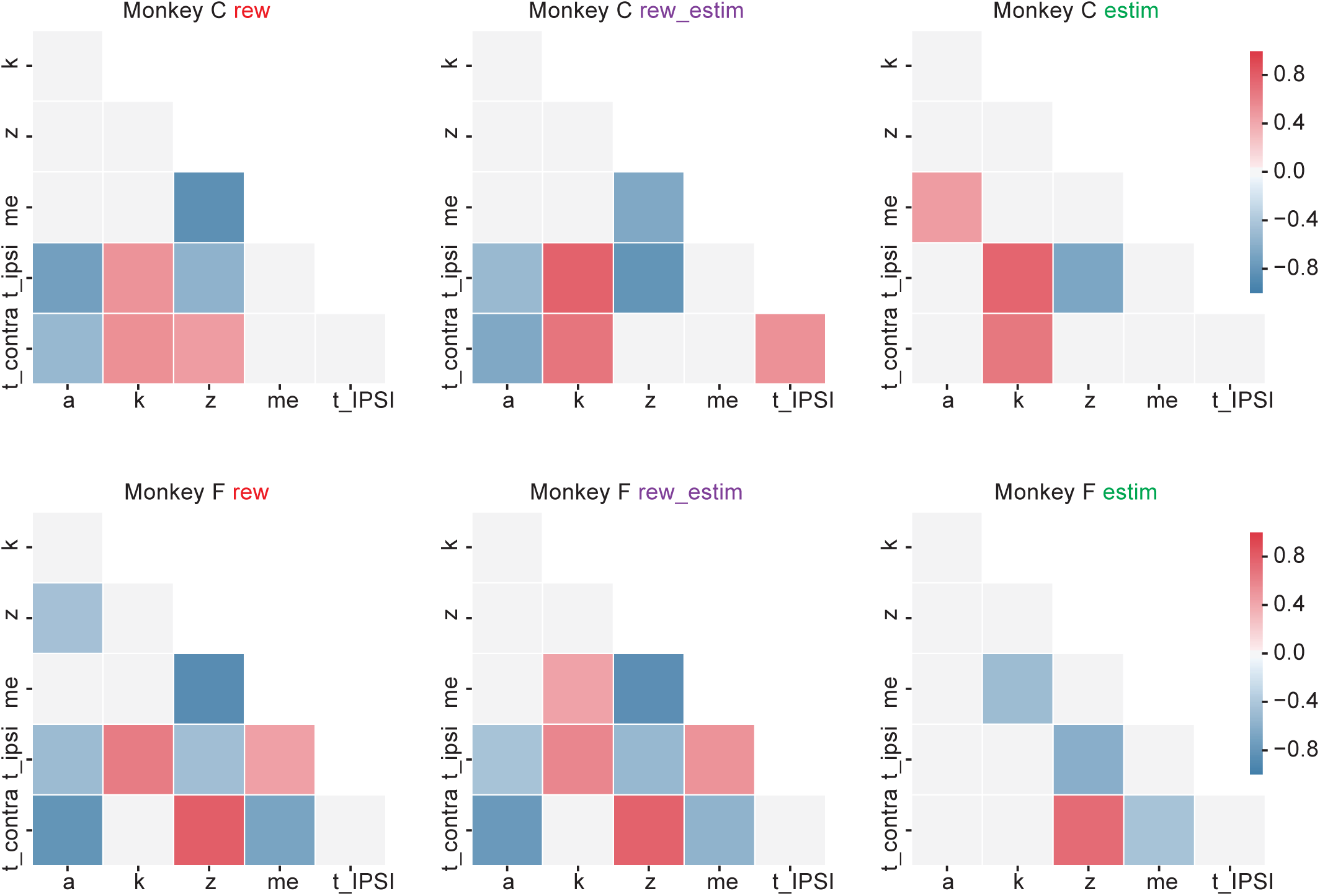
Pairwise correlation for the two monkeys. Same format as Fig. 4d, but showing data from the two monkeys separately (rows).

**Extended Data Figure 6.**
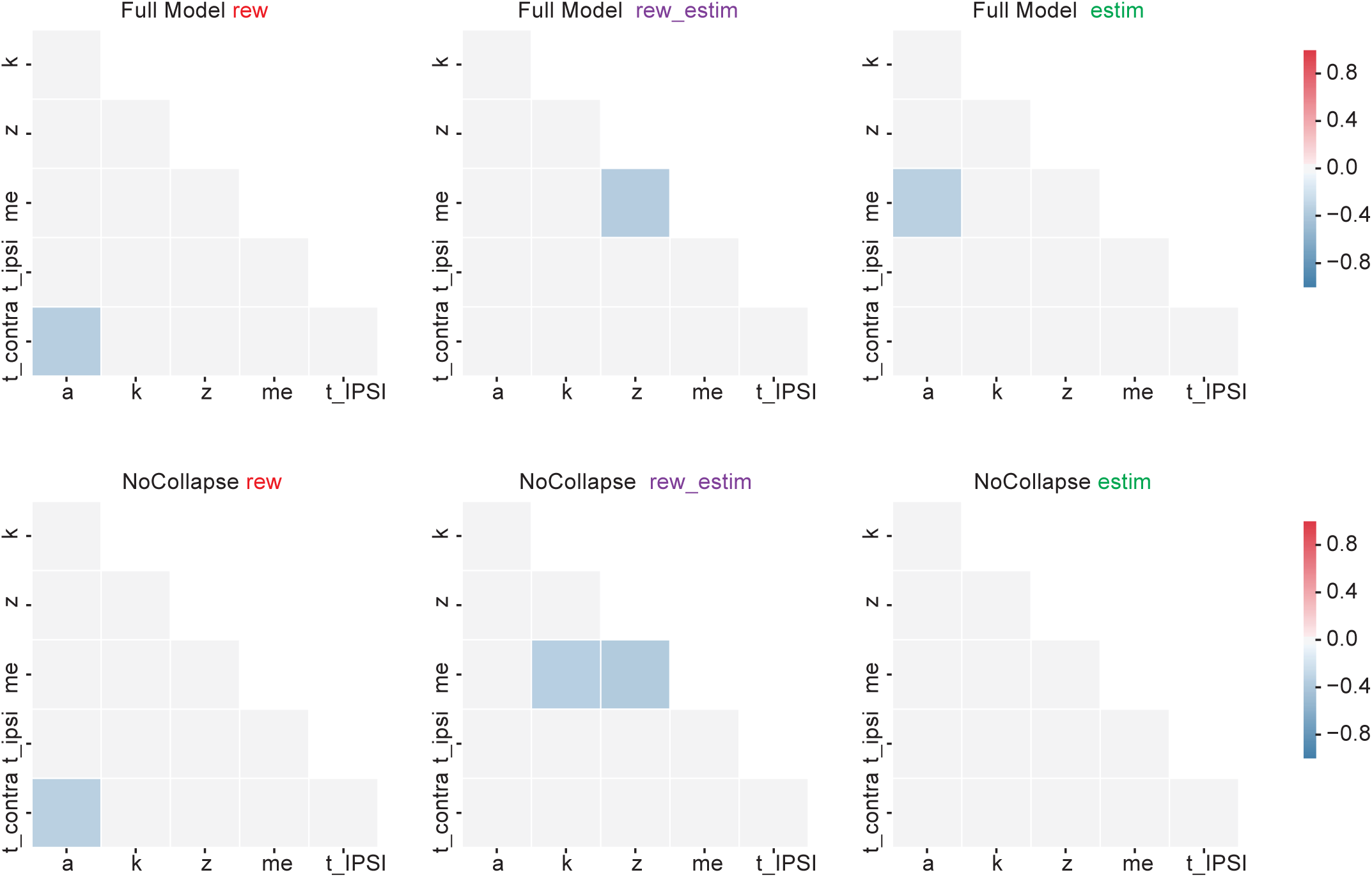
Pairwise correlation for simulated data fitted by the “Full” (top row) and “NoCollapse” (bottom row) models. Choice and RT data were simulated using the DDM with random combinations of parameter values resampled from model fits from all individual sessions. Same format as Fig. 4d. Note the overall absence of consistent relationships between pairs of best-fitting parameter values.

**Extended.**
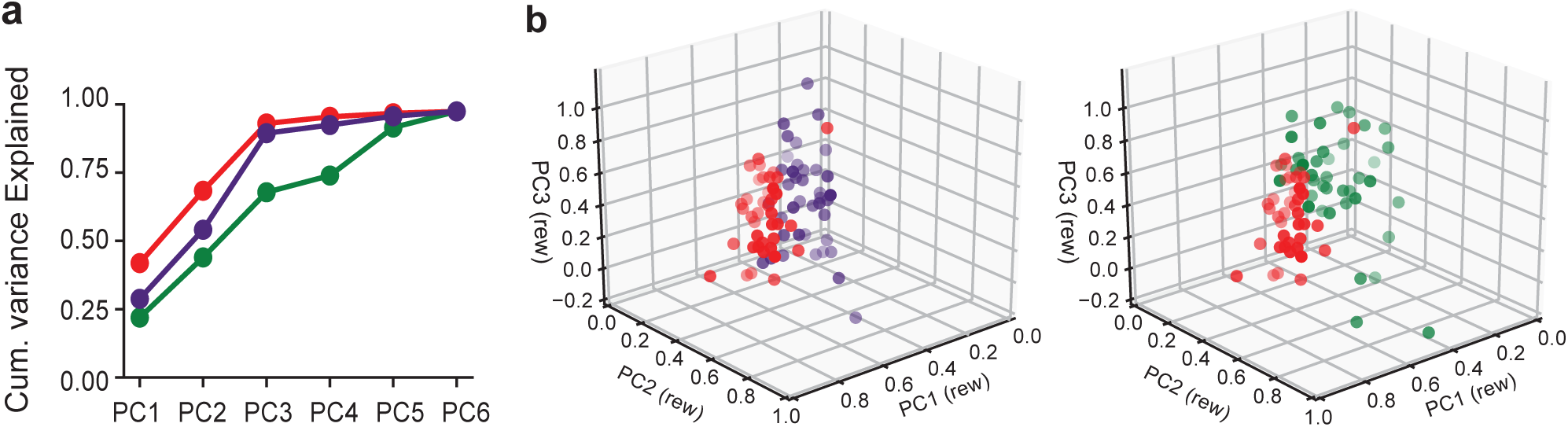
Data Figure 7. Principal component analysis. a) Cumulative fractions of variance of the session-specific values of best-fitting DDM parameters explained by principal components (PCs) of those parameters computed using best-fitting values of reward asymmetry-induced adjustments from non-microstimulation trials. Red: reward asymmetry-induced adjustments from non-microstimulation trials (“rew”), from which the PCs were extracted. Purple: reward context-dependent microstimulation effects (“rew-estim”). Green: reward context-independent microstimulation effects (“estim”). b) Projections of “rew” (red), “rew-estim” (purple) and “estim” (green) effects on DDM parameters in a 3D space defined by the first three principal components derived from the “rew” effects. The same “rew” data are plotted in both panels.

